# β-Carbonic anhydrase SlβCA4 is required for normal growth, carbon assimilation and stomatal plasticity in tomato

**DOI:** 10.64898/2026.10.21.759893

**Authors:** Orr Meyers, Mia Stern Tal, Jyoti Singh, Yuval Nagar, Iris Biton, Adi Yarran, Yaarit Kutsher, Yotam Zait, Ido Nir

## Abstract

β-Carbonic anhydrases (βCAs) catalyze the reversible interconversion of CO_2_ and bicarbonate and contribute to plant carbon metabolism and environmental responses. However, the physiological functions of individual βCA isoforms in crops remain poorly understood. Here, we investigated two chloroplast-localized βCAs, SlβCA1 and SlβCA4, in tomato (*Solanum lycopersicum*). We combined subcellular localization, CRISPR/Cas9 mutagenesis, grafting, leaf gas-exchange measurements, and controlled environmental treatments to examine their roles in growth, photosynthesis, and stomatal developmental plasticity. Both proteins localized to chloroplasts but their disruption resulted in markedly different phenotypes. While SlβCA1 mutants showed largely normal growth and photosynthesis, SlβCA4 mutants exhibited severe root and shoot growth defects. Grafting mutant shoots onto wild-type rootstocks partially restored vegetative growth, indicating that impaired root function contributes to the growth defects. SlβCA4 mutants also showed reduced CO_2_ assimilation and stomatal conductance, while electron transport remained largely unaffected. Elevated CO_2_ partially improved leaflet growth but did not restore plant height or root elongation. Furthermore, unlike wild-type plants, SlβCA4 mutants failed to increase stomatal index in response to elevated temperature. These findings reveal distinct physiological roles for chloroplast-localized βCAs in tomato and identify SlβCA4 as an important contributor to plant growth, photosynthesis, and temperature-dependent stomatal developmental plasticity.

## Introduction

Carbonic anhydrases (CAs) catalyze the reversible hydration of CO_2_ to bicarbonate (HCO3^−^) and protons, linking inorganic-carbon chemistry to plant metabolism and environmental responses. Because uncatalyzed interconversion can be slow relative to biological demand, the location and activity of CA isoforms may influence the balance of CO_2_ and HCO_3_^−^ in different tissues and therefore cellular inorganic carbon movements (Badger & Price, 1994; Ignatova *et al*., 2019; Momayyezi *et al*., 2020). In leaves, CA activity has been implicated in photosynthetic carbon assimilation and mesophyll conductance, but its physiological scope is broader: CA isoforms also participate in CO_2_-dependent stomatal responses, development, growth, and stress-related processes (Hu *et al*., 2010, 2021; Engineer *et al*., 2014; Hines *et al*., 2021). Thus, CA function cannot be inferred solely from a proposed role in CO_2_ diffusion to Rubisco.

Plant CAs comprise several protein families, including α-, β-, and γ-CAs, with individual isoforms differing in abundance, tissue expression, and subcellular localization (Momayyezi *et al*., 2020). βCAs are particularly abundant in C_3_ plants and occur in multiple cellular compartments, suggesting that individual isoforms perform distinct physiological functions. In Arabidopsis thaliana, six βCAs have been identified, with different expression patterns and subcellular localizations (Fabre *et al*., 2007; Momayyezi & Guy, 2017). Genetic studies have implicated βCA2, βCA4, and βCA5 in plant growth and development (DiMario *et al*., 2018; Weerasooriya *et al*., 2022), while βCA1, βCA4, and βCA6 have been associated with CO_2_-dependent responses, including the regulation of stomatal development (Hu *et al*., 2010; Engineer *et al*., 2014). Studies of the plastid-localized βCAs in Arabidopsis have further revealed an important distinction between their roles in plant growth and photosynthetic carbon assimilation. Loss of plastid CA activity can strongly impair Arabidopsis growth without substantially reducing photosynthetic CO_2_ assimilation, while the severe growth defect associated with loss of βCA5 can be alleviated by elevated CO_2_ (Hines *et al*., 2021; Weerasooriya *et al*., 2022, 2026). βCA1 has also been associated with carbon metabolism and physiological responses to changing environmental conditions in Arabidopsis (Rudenko *et al*., 2025). Together, these studies demonstrate that individual Arabidopsis βCA isoforms have distinct, but partially overlapping, roles in plant growth and carbon-related physiology.

In tomato (*Solanum lycopersicum*), four βCA genes have been identified: SlβCA1, SlβCA2, SlβCA3, and SlβCA4 (Feng *et al*., 2020; Lv *et al*., 2026). Their distinct expression patterns and predicted subcellular localizations suggest functional diversification among the tomato βCAs (Diamantopoulos *et al*., 2013; Feng *et al*., 2020). Recent studies have begun to define specific functions of SlβCA2 and SlβCA3, including roles in CO_2_-associated thermotolerance and basal immunity (Hu *et al*., 2021; Lv *et al*., 2026). In contrast, the cellular localization and physiological functions of SlβCA1 and SlβCA4 remain poorly understood. Phylogenetically, SlβCA1 groups closely with AtβCA1 and AtβCA2, whereas SlβCA4 is more closely related to AtβCA5 and AtβCA6 (Feng *et al*., 2020). Because these Arabidopsis homologs differ in both subcellular localization and physiological function, it remains unclear whether SlβCA1 and SlβCA4 perform overlapping or distinct roles in tomato.

Here, we investigated the subcellular localization and physiological functions of SlβCA1 and SlβCA4 in tomato. We first experimentally determined their cellular localization and then generated CRISPR/Cas9 mutant alleles to examine their contributions to plant growth and development. We further used grafting to assess the contribution of root function to the severe developmental phenotype associated with loss of SlβCA4, measured gas exchange to determine the effects of the mutations on photosynthetic CO_2_ assimilation, and tested whether elevated atmospheric CO_2_ could alleviate mutant growth defects. Together, these approaches reveal distinct functions for the two chloroplast-localized βCAs and identify SlβCA4 as an important regulator of tomato growth and carbon assimilation.

## Materials and Methods

### Growth conditions

Tomato plants (*Solanum lycopersicum* cv. M82) were grown in Conviron growth chambers (GEN2000, Conviron, Winnipeg, MB, Canada) set to a photoperiod of 16 hr light/8 hr dark and a light intensity of 300 µmol m^−2^ s^−1^. The temperature experiment was conducted under two conditions: 26°C with a relative humidity (RH) of 70%, or 34 °C with an RH of 80%, maintaining a constant VPD of ~1.03 kPa. The CO_2_ experiment was conducted in Aralab growth chambers (FitoClima 1200, Aralab, Rio de Mouro, Portugal) at CO_2_ concentrations of 390 or 800 ppm, a temperature of 26 °C, a relative humidity of 70%, and a photosynthetic photon flux density of 500 mol m^−2^ s^−1^. The photoperiod was the same as described above. For propagation, plants were grown in a greenhouse under natural day length conditions at 25–30 °C.

### Nicotiana benthamiana transformation

For tomato SlβCA marker lines for cellular localization analysis, transient expression in *Nicotiana benthamiana* was performed. The CDS of *SlβCA*s were first cloned into a level 0 MoClo part, then *35Sx2*_*pro*_*::SlβCA1-Venus, 35Sx2*_*pro*_*::slβca1-cr#5-Venus, 35Sx2*_*pro*_*::SlβCA1-exon1-Venus*, and *35Sx2*_*pro*_*::SlβCA4-Venus* were cloned using the Golden Gate system (Engler *et al*., 2014) into the binary vector pICH47742 with NOS terminator. The binary vectors were introduced into *Agrobacterium tumefaciens* strain GV3101 and infiltrated into mature tobacco leaves as described in Zhang et al. (2015). Plants were maintained for two days before tissue collection for Venus marker visualization using Stellaris confocal microscopy (STELLARIS 8, Leica Microsystems, Mannheim, Germany).

### Tomato CA1 and CA4 CRISPR/Cas9 mutagenesis, plant transformation, and selection of mutant alleles

To generate loss-of-function mutations in the SlβCA genes, three single-guide RNAs (sgRNAs) targeting distinct regions of the genomic sequence (Brooks *et al*., 2014). The sgRNAs were assembled into designated CRISPR/Cas9 binary vectors (pAGM4723, including zCas9 and NPTII) using the Golden Gate cloning system (Engler *et al*., 2014). The final vectors were introduced into *Agrobacterium tumefaciens* strain GV3101 by electroporation and transferred into M82 cotyledons using transformation and regeneration methods described by McCormick 1991 (McCormick, 1991). Kanamycin-resistant T0 transgenic plants were grown, and 10 to 15 independent lines were selected. For genotyping, genomic DNA was extracted, and each line was analyzed by PCR for the presence of zCas9 and characterized for specific mutations in *SlβCA1* and *SlβCA4* using gel electrophoresis and DNA sequencing. These lines were then grown over additional generations to isolate homozygous mutant lines, free from Cas9, for additional study.

### Light Microscopy sample prep and stomatal index counting

Cotyledons were harvested once plants had developed three visible true leaves. Cotyledons were cleared in 7:1 ethanol: acetic acid (4 h), rinsed in 1M KOH (30 min), and mounted in Hoyer’s solution. Images of the abaxial surface were taken at 20X using a light microscope (Rebel, ECHO, San Diego, CA, USA). Stomata were counted within a 0.4 mm^2^ leaf lamina region, avoiding vein areas. Cells were counted using FIJI (Schindelin *et al*., 2012) with the Cell Counter plugin. Results were plotted using GraphPad Prism. Stomatal index was calculated as [(no. of stomata) / (no. of stomata + no. of pavement cells)] * 100.

### Grafting

Seeds were sown in standard potting soil and maintained in a growth room under controlled temperature, humidity, and light conditions. Grafting was initiated when the seedling stems reached a diameter suitable for the grafting clips (~2 mm). Before the procedure, seedlings of *ca4-cr#1-1, ca4-cr#1-2, and* M82 were carefully uprooted and temporarily placed in water to photograph the intact whole plant, including the root system. For the grafting process, diagonal cuts (at an angle greater than 90 degrees) were made on the stems of both the mutant shoots and the M82 rootstocks. The cut surfaces were aligned and secured using specialized grafting clips designed for young seedlings. Thin wooden skewers were placed adjacent to the stems to provide structural support. Immediately following the procedure, the grafted plants were placed in a tray covered with a transparent plastic dome and sprayed with water to maintain a high-humidity environment. The plants were kept under these conditions until a complete and stable junction formed between the scion and the rootstock, at which point the fully healed plants were photographed again before being transferred to the greenhouse for additional experiments.

### Gas-exchange measurements

To quantify differences in physiological parameters among *S. lycopersicum* genotypes, net CO_2_ assimilation rate (A_n_), stomatal conductance to water vapor (g_sw_), and electron transport rate (ETR) were measured using a LI-6800F portable photosynthesis system (LI-COR Biosciences, Lincoln, NE, USA) equipped with a 2 cm^2^ leaf chamber. Measurements were performed on the youngest fully expanded leaf of each plant, ensuring a comparable developmental stage across genotypes. Measurements were conducted at areference CO_2_ concentration of 420 µmol CO_2_ mol^−1^ air, and a flow rate of 500 µmol air s^−1^. Relative humidity inside the leaf chamber was maintained between 50% and 60%.

### Gene identifiers

S. lycopersicum: SlCA1 (*Solyc02g086820*); SlCA4 (*Solyc09g010970*)

### Quantification and statistical analyses

Statistical analyses were performed using GraphPad Prism version 8.0.2 (GraphPad Software, San Diego, California, USA), and RStudio (R Development Core Team., 2020). Statistical parameters for each analysis, including sample sizes (n), are indicated in the figure legends. To assess the main effects of genotype and treatment, as well as their interaction, data were analyzed using a Two-Way Analysis of Variance (ANOVA), followed by Tukey’s multiple comparisons test conducted for post-hoc analysis, or Games-Howell post-hoc tests (Games & Howell, 1976) using the games_howell_test function from the rstatix package (Kassambara, 2021). In the figures, different letters indicate groups of samples with no statistically significant difference between their means (p > 0.05).

## Results

### Chloroplast localization of SlβCA1 and SlβCA4

Previews studies show that the tomato SlβCA1 cluster closely with the *Arabidopsis* AtβCA1, while SlβCA4 cluster closely with AtβCA5 and AtβCA6, suggesting that these two tomato SlβCAs are plastid localized (Fabre *et al*., 2007; Weerasooriya *et al*., 2022). To examine their subcellular localization, we first performed an *in silico* analysis of the SlβCA1 and SlβCA4 protein sequences using the TargetP-2.0 platform (Almagro Armenteros *et al*., 2019); Table S1). This analysis predicted that SlβCA1 contains a functional chloroplast transit peptide, whereas SlβCA4 was predicted to have partial mitochondrial localization. To experimentally their subcellular localization, we transiently expressed SlβCA1 and SlβCA4 fused to a Venus fluorescent reporter under the constitutive 35S promoter, (35S::SlβCA1-Venus and 35S::SlβCA4-Venus) in *Nicotiana benthamiana*. Confocal microscopy confirmed that both SlβCA1 and SlβCA4 localize to chloroplasts (Figures 1a and 1d), despite the predicted partial mitochondrial localization of SlβCA4 (Table S1).

**Figure 1.**
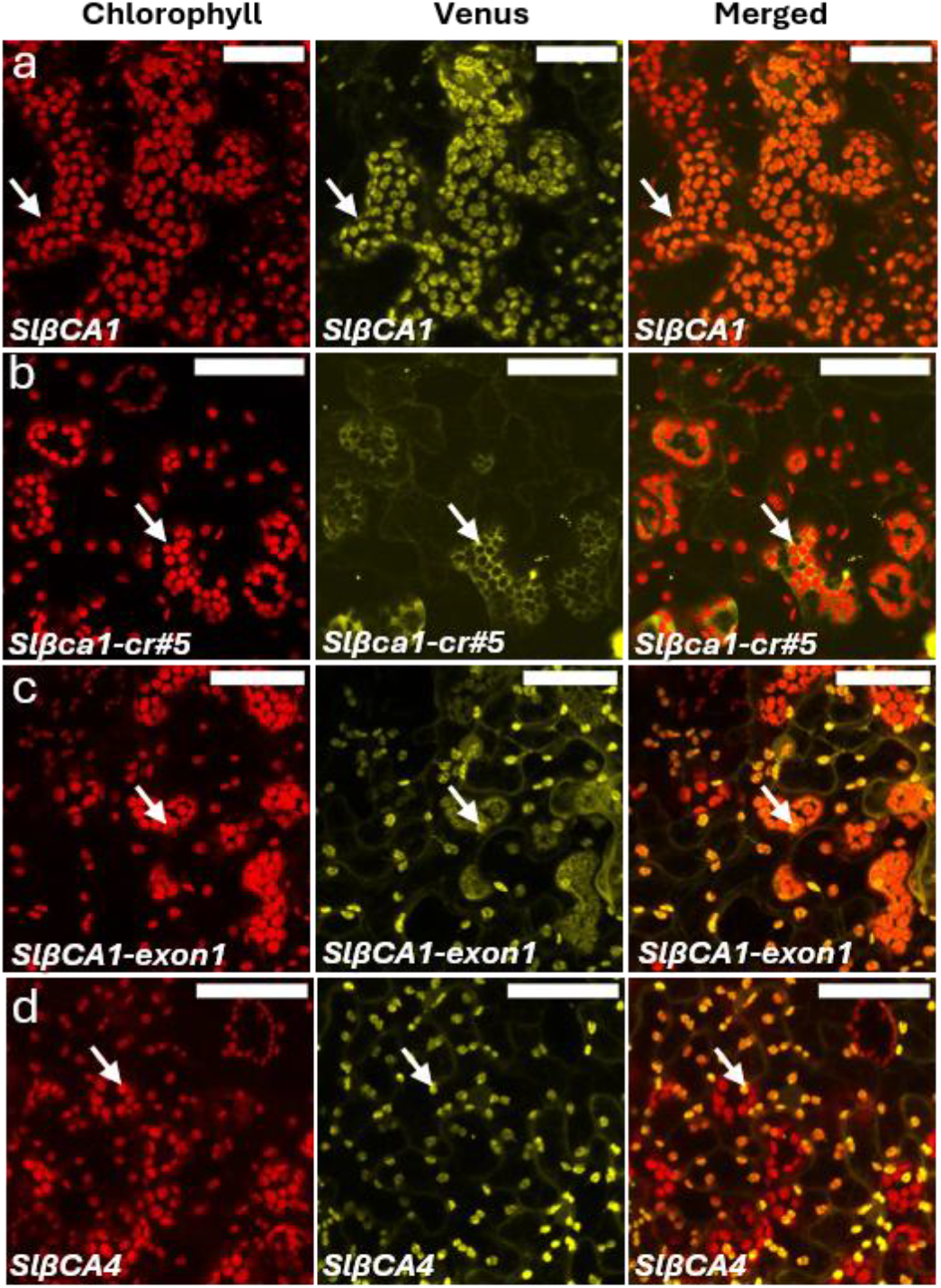
Subcellular localization of SlβCA variants in *N. benthamiana*. Confocal images (Maximum Intensity Projections) of transiently expressed Venus-fused constructs 48h post-agroinfiltration. (a) Wild-type *35S::SlβCA1-Venus*. (b) Mutant *35S::Slβca1-cr#5-Venus*. (c) *35S::SlβCA1-exon1-Venus*. (d) Wild-type *35S::SlβCA4-Venus*. Columns from left to right: Chlorophyll autofluorescence (red), Venus fluorescence (yellow), and merged images. White dashed lines indicate mesophyll cell boundaries. Scale bars = 50 µm.

To investigate the physiological functions of these two chloroplast-localized βCAs, we next used CRISPR/Cas9 genome editing to generate mutations in SlβCA1 and SlβCA4. Three single-guide RNAs (sgRNAs) were designed to target distinct regions of each gene (Brooks *et al*., 2014). We selected 10 to 15 independent lines per gene, characterized the mutations by PCR and DNA sequencing, and propagated them to identify homozygous mutant plants. For *SlβCA1*, we focused on two lines: *ca1-cr#5*, which contains a large deletion eliminating exon 1 but retains an alternative start codon (ATG) in exon 2, and *ca1-cr#7*, which contains a structural inversion (Figure 2a). Although neither SlβCA1 mutant displayed an obvious developmental phenotype (Figure 2c), the nature of the *ca1-cr#5* mutation prompted us to examine whether it affected SlβCA1 subcellular localization. Because *ca1-cr#5* lacks exon 1 but retains an alternative ATG in exon 2, we predicted that this allele could produce a truncated SlβCA1 protein lacking the N-terminal chloroplast transit peptide. TargetP-2.0 analysis of the predicted *ca1-cr#5* protein supported this prediction (Table S1). We therefore transiently expressed the *ca1-cr#5* protein fused to Venus in *N. benthamiana*. In contrast to wild-type SlβCA1, the *ca1-cr#5* failed to accumulate in chloroplasts and instead localized to other subcellular compartments (Figure 1b). Moreover, expression only SlβCA1 exon 1 fused to Venus was sufficient to target the reporter to chloroplasts (Figure 1c), demonstrating that exon 1 contains the functional chloroplast-targeting sequence of SlβCA1. Thus, although *ca1-cr#5* plants did not exhibit an obvious developmental phenotype, the mutation disrupted SlβCA1 chloroplast localization.

**Figure 2.**
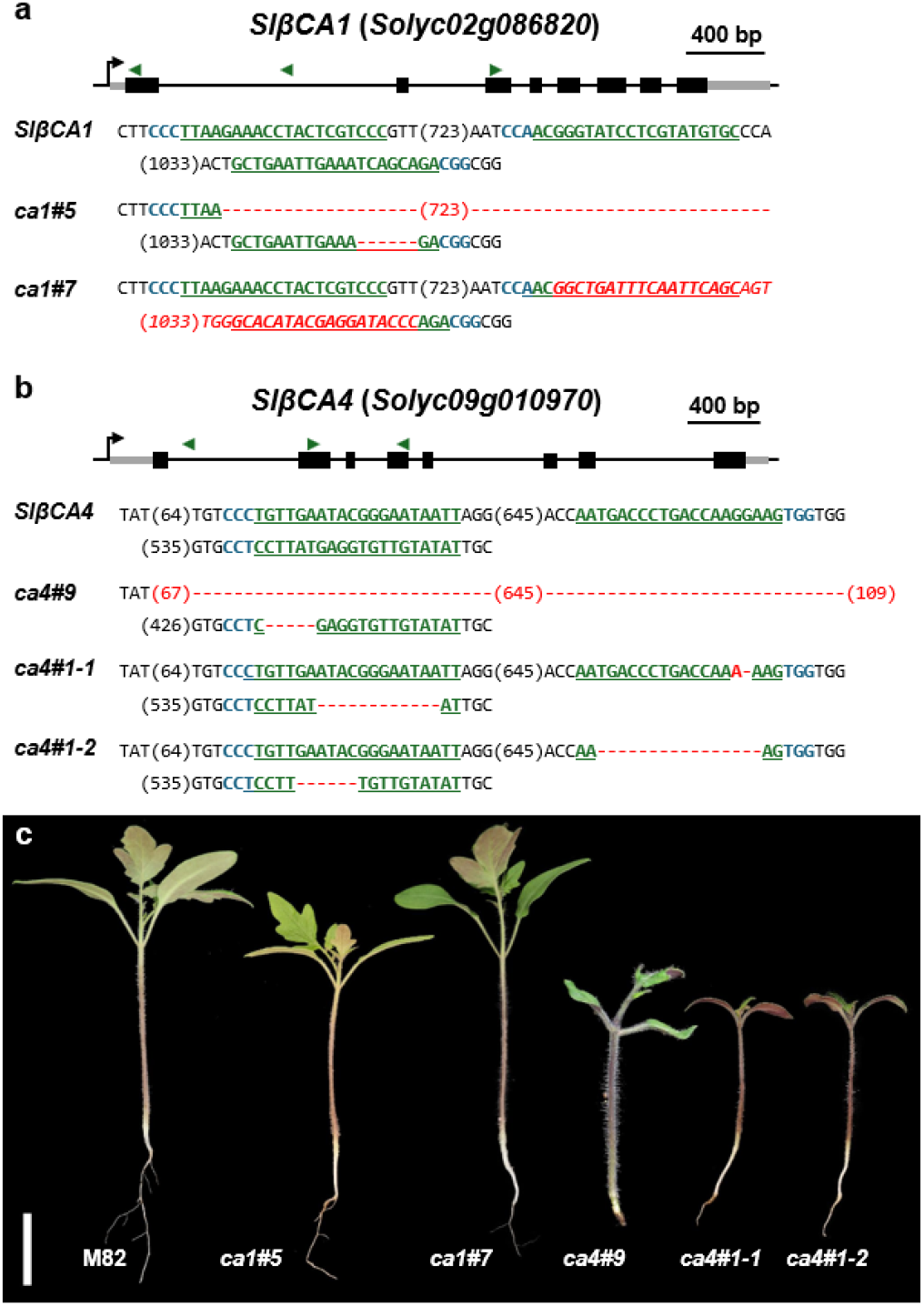
Generation of SlβCA1 and SlβCA4 CRISPR mutants reveals a distinct developmental phenotype in SlβCA4 mutants. (a,b) Gene structures and molecular characterization of CRISPR/Cas9-induced mutations in *SlβCA1* (a) and *SlβCA4* (b). Exons and untranslated regions (UTRs) are represented by black and grey boxes, respectively, while introns are shown as black lines. Bent arrows indicate the transcription start sites, and green triangles mark the sgRNA target locations. Sequence alignments of the wild-type and the generated CRISPR mutant alleles are shown below each gene model. The sgRNA target sequences are underlined in green, and the PAM sequences are highlighted in blue. Red dashes indicate deletions, and red text indicates nucleotide insertions or structural changes (e.g., inversions). Numbers in parentheses represent the length of unshown intervening sequences or large deletions in base pairs. (c) Representative seedling phenotypes of M82 and T1 mutant lines of *slca1-cr#5* (*ca1#5*), *slca1-cr#7* (*ca1#7*), *slca4-cr#9* (*ca1#9*), *slca4-cr#1-1* (*ca4#1-1*), and *slca4-cr#1-2* (*ca1#1-2*), Scale bar = 2 cm. Due to low germination rates, *ca4#9* was grown independently and photographed at an older developmental stage compared to the other plants.

In contrast to SlβCA1, generating viable *SlβCA4* mutants was considerably more challenging because mutations in this gene severely impaired plant development. Among the 26 T0 lines generated, 3 were homozygous for deletion ranging from 2 bp to 650 bp and displayed severe developmental defects, most notably a dramatic reduction in root and shoot growth (Figure S1). Most of the homozygous mutants could not be propagated due to impaired reproductive development. Two lines, *ca4-cr#1* and *ca4-cr#9*, carried heterozygous mutations and were able to produce seeds, allowing further characterization of the mutant alleles (Figure 2b and c). In subsequent generations, *ca4-cr#1* segregated into two alleles, *ca4-cr#1-1* and *ca4-cr#1-2*, from which homozygous mutant plants were obtained. Similarly, *ca4-cr#9* produced homozygous plants carrying a large deletion (Figure 2b). Phenotypic characterization of these independent SlβCA4 alleles consistently revealed severe developmental defects, including extreme dwarfism and strongly impaired root growth (Figure 2c). These phenotypes indicate that SlβCA4 is required for normal tomato growth and development, but do not distinguish whether its function is required in the root, the shoot, or both. To address this question, we next performed grafting experiments between wild-type and SlβCA4 mutant plants.

### Wild-type rootstocks partially rescue the growth defects of slca4-cr#1 mutant

The severe root and shoot growth defects of the SlβCA4 mutants raised the question of whether SlβCA4 function is required independently in both tissues or whether the impaired shoot growth could, at least in part, result from the defective root system. To address this, we grafted *ca4-cr#1* mutant shoots onto M82 rootstocks, and M82 shoots onto M82 rootstocks as control (Figures 3b and S2), to determine whether a functional root system could improve mutant shoot growth. Following a recovery period under high-humidity conditions to promote tissue healing and acclimatization, the grafted plants successfully established stable graft unions (Figure 3b). Notably, *ca4-cr#1* shoots grafted onto M82 rootstocks exhibited substantial vegetative growth and developed into mature plants (Figure S3), with improved growth compared with non-grafted mutants (Figure 3c). These results indicate that a functional wild-type root system can partially alleviate the shoot growth defects caused by loss of SlβCA4 function, suggesting that the severe shoot phenotype is, at least in part, associated with impaired root function. However, grafting did not fully restore normal shoot development, indicating that the SlβCA4 phenotype cannot be explained solely by impaired root function.

**Figure 3.**
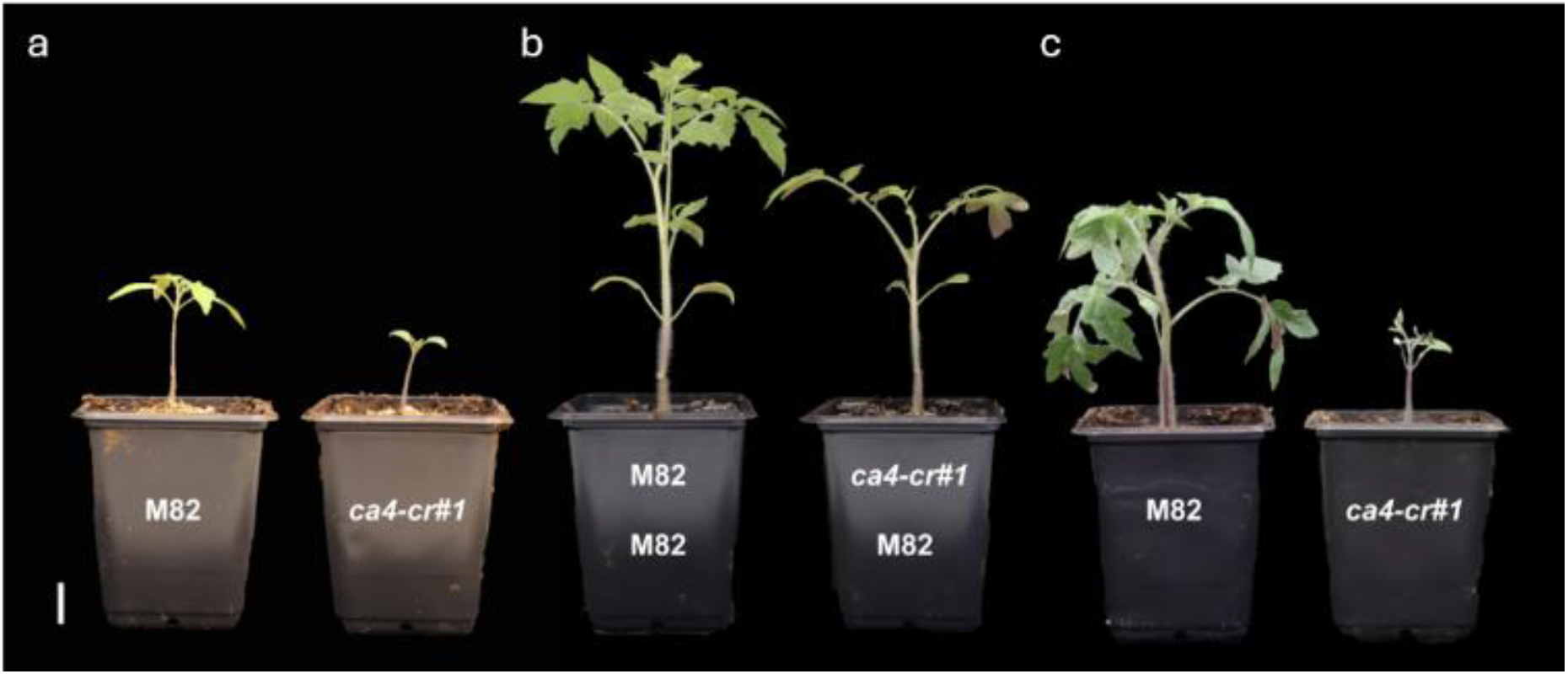
Improved growth of *SlβCA4* mutants by grafting onto M82 rootstocks. (a, c) Non-grafted plants showing the severe dwarf phenotype of *slβca4* mutants compared with the normally developing M82 wild type. Plants are approximately 3 weeks old in (a) and 6 weeks old in (c). (b) Grafted plants, approximately 6 weeks old, demonstrate a partial morphological rescue of the mutant shoots compared with a self-grafted M82 control. Scale bar = 2 cm.

### Loss of SlβCA4, but not SlβCA1, reduces photosynthetic CO_2_ assimilation

Given the chloroplast localization of SlβCA4 and the established role of βCAs in facilitating CO_2_ availability for photosynthesis (Badger & Price, 1994), we next asked whether loss of SlβCA4 affects leaf photosynthetic performance independently of its severe root phenotype. We therefore measured leaf gas exchange and electron transport in *ca4-cr#1* mutant shoot grafted onto M82 rootstocks and compared them with M82 shoots grafted onto M82 rootstocks. We also analyzed the SlβCA1 mutant lines, which develop normally on their own roots, to determine whether disruption of the two chloroplast-localized βCAs has similar effects on photosynthetic function.

Marked differences between the *slca4-cr#1* lines and M82 were evident in gas-exchange parameters measured at an ambient CO_2_ concentration of 420 ppm (Figure 4a-c). Compared with M82, net CO_2_ assimilation (A) was reduced by 46.1% in *slca4-cr#1* _young (9 weeks) and by 12.7% in *slca4-cr#1*_mature (14 weeks), while stomatal conductance (*g*_*sw*_) decreased by 55.5% and 40.9%, respectively. In contrast, electron transport rate (ETR) was reduced by 16.0% in *slca4-cr#1*_young and by 0.2% in *slca4-cr#1* _mature, and these differences were not statistically significant. Thus, the reduction in photosynthetic CO_2_ assimilation in the *ca4* lines was accompanied by a pronounced decrease in stomatal conductance, whereas electron transport was largely maintained. In contrast to the *slca4-cr#1* line, the *slca1*-cr#5 and *slca1*-cr#7 lines exhibited no significant reductions in A, *g*_*sw*_, or ETR relative to M82 (Figure 4d-f). Both A and gsw were numerically similar to or higher than those of M82 in *slca1-cr#5* and *slca1-cr#7*, while ETR showed only minor, non-significant variation, with a 4.3% decrease in *slca1-cr#5* and a 4.8% increase in *slca1-cr#7*. These results indicate that disruption of SlβCA4, but not SlβCA1, was associated with reduced photosynthetic performance under ambient CO_2_ conditions.

**Figure 4.**
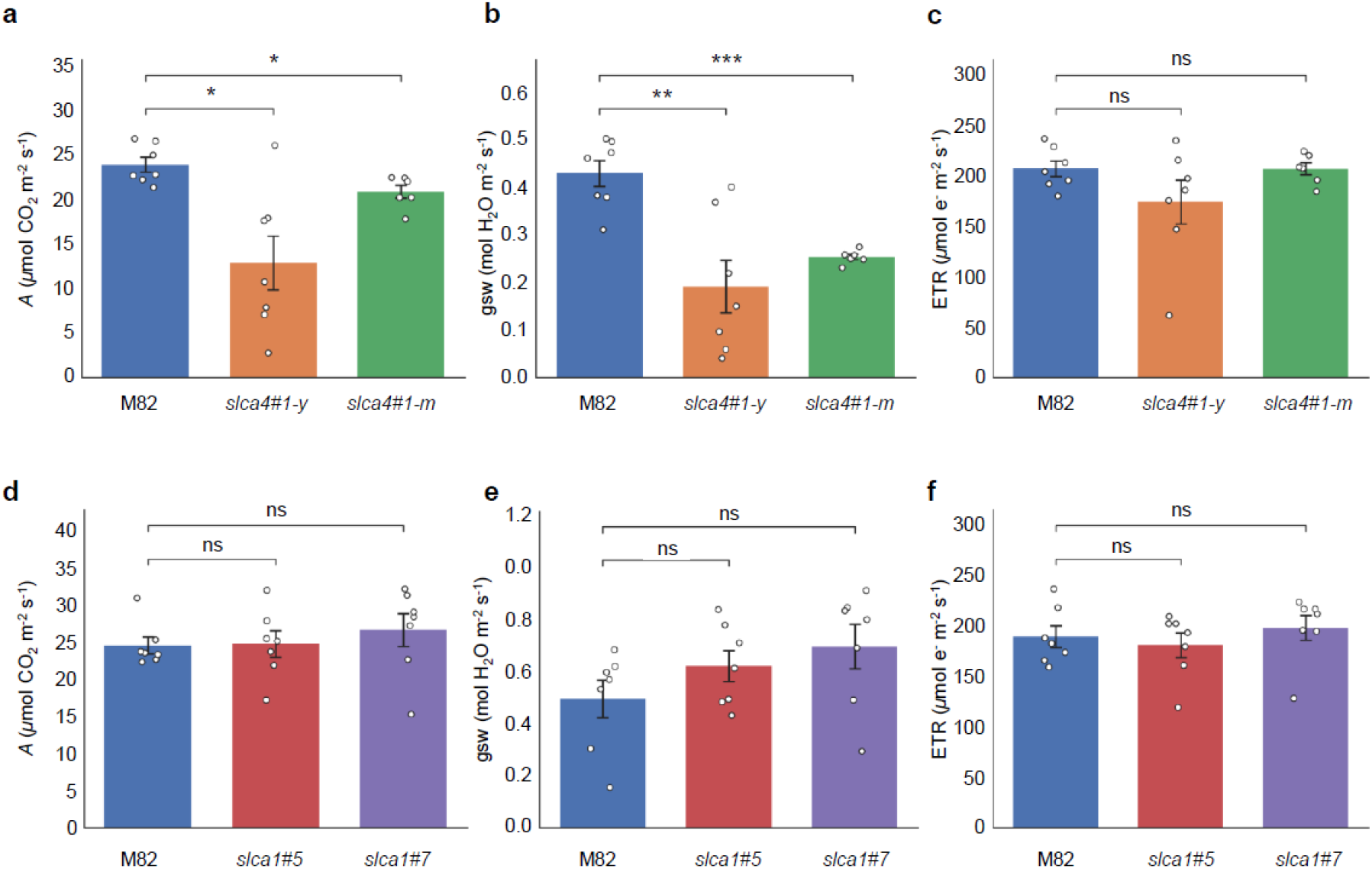
Photosynthetic performance of *slca4* and *slca1* mutant lines under ambient CO_2_ conditions. Net CO_2_ assimilation rate (A; µmol CO_2_ m^−2^ s^−1^), stomatal conductance to water vapor (gsw; mol H_2_O m^−2^ s^−1^), and electron transport rate (ETR; µmol e^−^ m^−2^ s^−1^) of *slca4* and *slca1* mutant lines. (a-c) A, gsw, and ETR, respectively, in M82, slca4#1-y, and slca4#1-m. (d-f) A, gsw, and ETR, respectively, in M82, slca1#5, and slca1#7. Bars represent means ± SE, and individual points represent biological replicates. For (a-c), n = 7 for M82 and slca4#1-y and n = 6 for slca4#1-m; for (d-f), n = 7 for each genotype. Statistical significance was assessed using unpaired two-tailed Welch’s t-tests comparing each mutant line with M82. Asterisks indicate statistical significance (P < 0.05 (*), P < 0.01 (**), and P < 0.001 (***)); ns indicates no statistically significant difference.

### Elevated CO_2_ partially rescues the growth defect of SlβCA4 mutants

The reduced CO_2_ assimilation observed in *<u>ca4-cr#1</u>* raised the possibility that limited carbon assimilation contributes to its severe growth phenotype. We therefore asked whether increasing atmospheric CO_2_ availability could alleviate the developmental defects caused by loss of SlβCA4. To test this, we compared the growth of M82, ca1-cr#5, and *ca4-cr#1* plants under 390 ppm and elevated (800 ppm) CO_2_ conditions. Under ambient CO_2_, ca1-cr#5 plants showed growth comparable to M82, whereas *ca4-cr#1* plants exhibited severely stunted growth, with significantly reduced plant height and leaf area (Figure 5a-c). Elevated CO_2_ significantly increased plant height in both M82 and *ca1-cr#5*, whereas no significant increase in plant height was observed in *ca4-cr#1* (Figure 5a and b). Leaflet area showed a different response. Elevated CO_2_ reduced leaf area in M82 and ca1-cr#5, whereas it significantly increased leaf area in *ca4-cr#1* (Figure 5c). Despite this increase, the leaf area of *ca4-cr#1* remained significantly lower than that of M82 and *ca1-cr#5* under elevated CO_2_.

**Figure 5.**
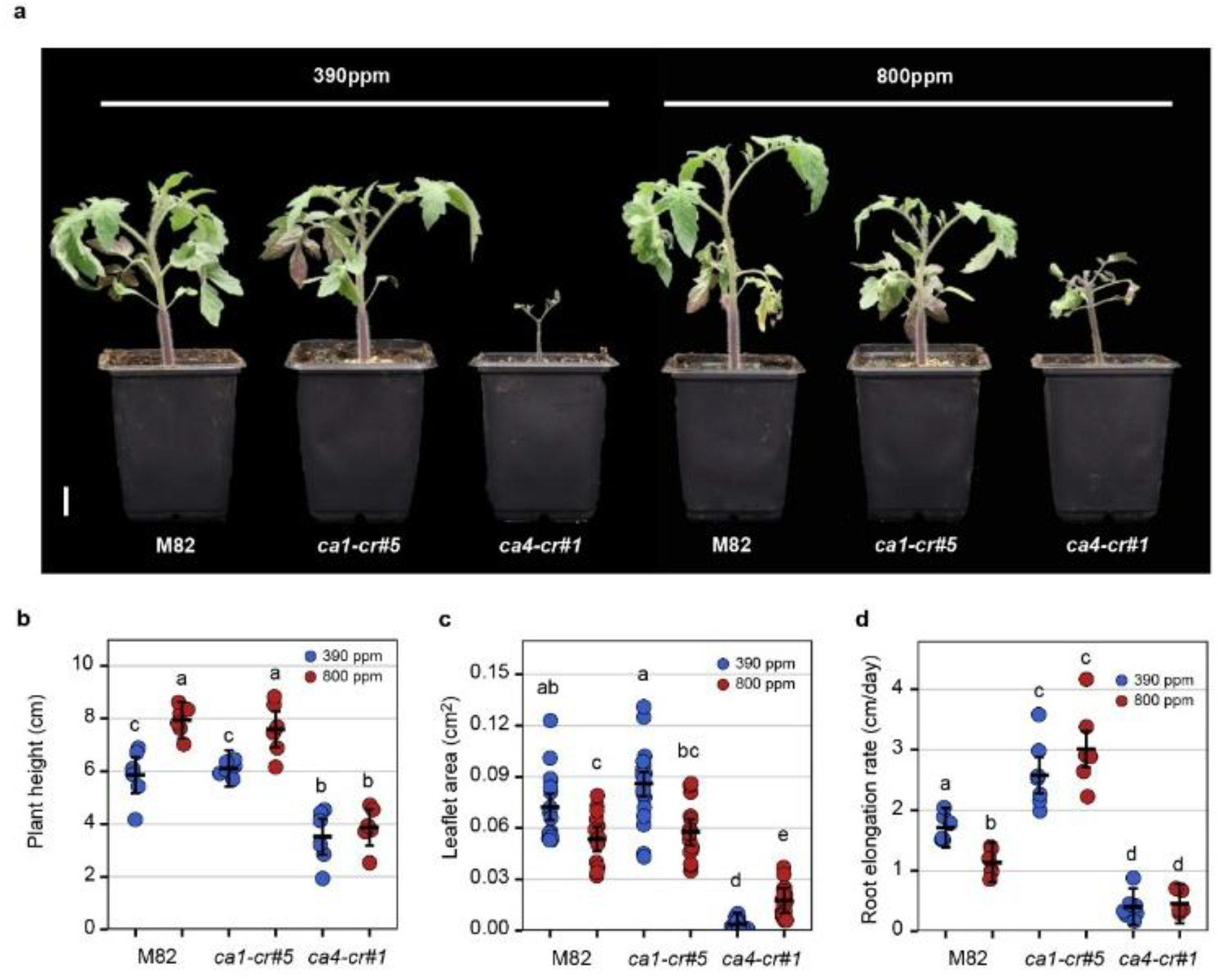
Phenotypic response of wild-type and mutant lines to elevated CO_2_ concentrations. (a) Representative images of tomato wild-type (M82) and SlβCA mutants (*ca1-cr#5* and *ca4-cr#1*) grown under ambient (390 ppm) and elevated (800 ppm) CO_2_ conditions. (b) Plant height, (c) leaf area, and (d) root elongation rate of the indicated genotypes under the respective CO_2_ treatments. Data are expressed as mean ± SD, with individual points representing biological replicates in (a) n = 6, in (b) n = 16-18 leaflet from 6 individual plants, and (d) n = 5-7 individual plants. Different letters indicate statistically significant differences among groups, as determined by two-way ANOVA followed by Tukey’s multiple comparisons test (P < 0.05).

We next examined whether elevated CO_2_ also affected root growth. Under ambient CO_2_, the root elongation rate of *ca4-cr#1* was strongly reduced compared with M82 and *ca1-cr#5* (Figure 5d). Elevated CO_2_ did not significantly improve root elongation in *ca4-cr#1*, which remained markedly lower than in the other genotypes. In contrast, elevated CO_2_ reduced root elongation in M82, while ca1-cr#5 maintained a substantially higher root elongation rate under both CO_2_ conditions (Figure 5d).

Thus, elevated CO_2_ partially alleviated the growth defect of *ca4-cr#1*, as reflected by the increase in leaflet area, but did not significantly restore plant height or root elongation. These results are consistent with reduced carbon assimilation contributing to aspects of the *ca4-cr#1* growth phenotype, while the persistence of the severe root-growth defect under elevated CO_2_ suggests that increased external CO_2_ availability is insufficient to compensate for the loss of SlβCA4 function.

### Stomatal developmental responses of SlβCA mutants to elevated temperature

Having established that the developmental phenotypes of *SlβCA* mutants are closely linked to carbon availability, we shifted our focus to the primary gateway for carbon entry: the stomata. Since previews works shown that the Arabidopsis AtβCA’s involve in the regulation of stomatal development and function in response to environmental signals (Hu *et al*., 2010; Engineer *et al*., 2014), we hypothesized that SlβCA1 and SlβCA4 might facilitate similar developmental plasticity in tomato. To test this, we conducted experiments in controlled climate chambers to evaluate stomatal responses to temperature changes. We grew control M82, together with *ca1-cr#5*, and *ca4-cr#1* mutant plants under standard (26°C) and elevated (34°C) temperatures and quantified the stomatal index (SI) to measure the developmental plasticity in these lines. In the M82 plants, raising the temperature from 26°C to 34°C induced a significant increase in SI. The *ca1-cr#5* mutant maintained a near-normal response capacity. Although its SI under 26°C conditions was slightly elevated, hiding the statistical significance of its internal temperature induction, it successfully reached maximal SI levels at 34°C that were statistically identical to those of the wild-type (Figure 6a). To determine the role of SlβCA4 in this dynamic, we similarly analyzed the *ca4-cr#1* mutant. Unlike *ca1-cr#5*, the *ca4-cr#1* plants displayed a loss of temperature responsiveness, showing no significant alteration in SI at 34°C (Figure 6b). These findings suggest that while SlβCA1 may play a minor role in modulating baseline stomatal development, SlβCA4 is strictly required for the temperature-induced developmental plasticity of the stomata.

**Figure 6.**
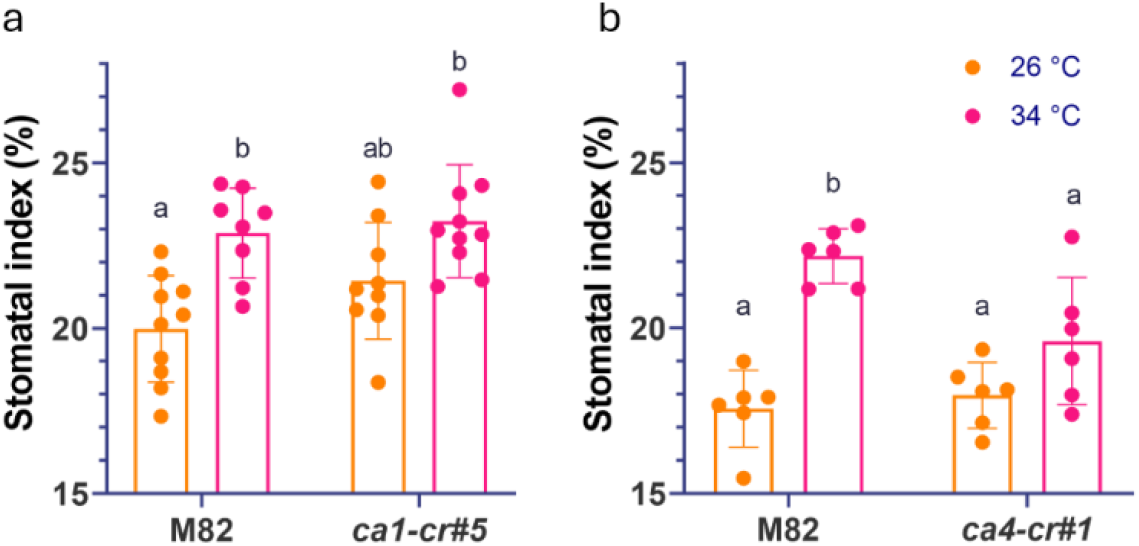
Stomatal index (SI) responses to elevated temperature in SlβCA mutants. Stomatal index of *ca1-cr#5* (a) and *ca4-cr#1* (b) compared to the M82 wild-type at 26°C and 34°C. Dots represent the mean SI per cotyledon (from 6 independent images, n = 6–10). Data are expressed as mean ± SD. Significance was determined by a Two-Way ANOVA followed by Tukey’s multiple comparisons test. Different letters indicate statistically significant differences between means (p < 0.05). SI = [stomata / (stomata + pavement cells)] × 100.

## Discussion

Whereas βCA enzymes have been studied extensively in *Arabidopsis* (Badger & Price, 1994; Engineer *et al*., 2014; Weerasooriya *et al*., 2022, 2026), their functions in crop species such as tomato remain partly characterized. In this study, we characterized the function of two tomato isoforms, SlβCA1 and SlβCA4. Initial molecular characterization of CRISPR/Cas9-edited lines revealed diverse mutation profiles in both *SlβCA1* and *SlβCA4*, ranging from small deletions of a few nucleotides (e.g., *slca4#1*) to large multi-base deletions (e.g., *slca1#5* and *slca4#9*) and inversions (e.g., *slca1#7*; Figure 1a and b). Crucially, regardless of the mutation type or size, all *slca4* homozygous mutant line consistently exhibited an exceptionally severe phenotype characterized by extreme dwarfism and stunted development. In contrast, *slca1* homozygous mutants displayed no discernible visual phenotype compared to the M82 control plants (Figure 1c). The lethality and severe developmental defects of *slca4* plants presented significant challenges in maintaining these lines, as most tissue-cultured regenerates T0 plants failed to survive the transition to the greenhouse or produce viable seeds. Consequently, we focused our downstream physiological analyses on the *slca4#1* line, which harbors two highly similar independent mutations in the same line (Figure 1b and c), alongside the homozygous *slca1#5* line. Notably, the dramatic developmental impairment observed in *slca4* closely resembles the severe phenotypes reported for mutants of its homologs, *AtβCA5* in *Arabidopsis* and *NtβCA5* in tobacco (Hines *et al*., 2021; Weerasooriya *et al*., 2022, 2026).

To further clarify the functional differences among these isoforms, we examined their subcellular localization. Consistent with in silico predictions (Feng *et al*., 2020) and the localization of their *Arabidopsis* and tobacco homologs (Hines *et al*., 2021; Weerasooriya *et al*., 2026), transient expression assays in tobacco leaves confirmed that both SlβCA1 and SlβCA4 localize to the chloroplast (Figure 1). Furthermore, our data suggest that the deletion in exon 1 of the c*a1#5* mutant reveals an alternative translation start codon (ATG) at the beginning of exon 2. This alternative translation produced a shortened protein that failed to enter the chloroplast, instead accumulating in cell cytosol while remaining excluded from chloroplasts (Figure 1b). An additional transient assay using only the exon 1 sequence confirmed its function as a chloroplast transit peptide (Figure 1c). Although this shortened, cytosolic protein in the *ca1#5* line might retain some enzymatic activity, this alternative translation cannot fully explain the absence of a severe phenotype in this gene. Crucially, all ca1 mutant lines, including those with mutations downstream of exon 1 (e.g., ca1#7), exhibited similarly mild phenotypes (Figure 2a and c). Therefore, we propose that the mild phenotype of ca1 mutants is primarily due to functional redundancy. It is likely that other CA isoforms, potentially including SlβCA4 itself, can effectively compensate for the loss of SlβCA1, whereas SlβCA4 performs an essential, non-redundant role for normal plant development. Our observation that the *slca1* mutant lines showed no obvious developmental phenotype and no significant reduction in steady-state An, gsw, or ETR under the conditions tested is consistent with recent evidence that the major chloroplast βCA1 isoform in tomato is not required primarily for direct CO_2_ supply to Rubisco. Wang et al. (2026) showed that tomato chloroplast βCA1 instead contributes to the coordination of CO_2_ availability and irradiance by buffering light-induced changes in stromal pH and thereby modulating proton-motive-force partitioning and PSII performance (Wang *et al*., 2026).

Characterizing the physiological role of SlβCA4 was initially delayed by the severely limited availability of viable T1 seeds. Attempts to propagate the *slca4* mutants through crosses with wild-type M82 plants resulted in fruits containing few viable seeds, and the resulting T1 seedlings failed to develop normally or reach maturity (Figures 2c and 3c). Ultimately, we were able to significantly improve the vegetative growth of the *slca4* mutants and achieve reproductive maturity by grafting *slca4* mutant scions onto M82 rootstocks (Figures 3, S2 and S3). We speculate that because *SlβCA4* is ubiquitously expressed across multiple tissues, including the root system (Feng *et al*., 2020), grafting onto a M82 rootstock partially compensates for the loss of the enzyme in the roots. Consequently, supported by healthy M82 roots, the mutant shoot can sustain itself by relying primarily on the intact SlβCA1, or other SlβCA enzymes, in the leaves. While we utilized grafting mainly as a methodological tool to recover functional seeds, these observations highlight the systemic importance of functional roots in mitigating the *slca4* mutants developmental arrest. Importantly, this grafting approach proved to be a successful strategy to bypass the severe barrenness and developmental arrest of the *slca4* mutants. By achieving reproductive maturity on M82 rootstocks, we were finally able to produce viable fruits from recent crosses with *SlβCA4* overexpressing lines. Ultimately, this methodological workaround successfully recovered functional seeds, ensuring the conservation and continuation of the *slca4* mutant lines.

Finally, applying the limited remaining seeds of the *ca4#1* line alongside the homozygous *ca1#5* line, we assessed the mutants’ responses to varying environmental CO_2_ concentrations. Because chloroplast stroma becomes alkaline during light exposure, inorganic carbon predominantly exists as HCO_3_^−^ (Rudenko *et al*., 2025). We hypothesized that disruption of SlβCA4 may alter inorganic-carbon homeostasis in a manner that limits growth and photosynthetic performance. However, recent evidence for tomato chloroplast βCA1 indicates that chloroplast βCAs can also regulate photosynthetic function through stromal-pH buffering and CO_2_ light coordination rather than through direct provision of CO_2_ to Rubisco (Wang *et al*., 2026).

Thus, our experiments were designed to test whether elevated external CO_2_ could alleviate the *slca4* phenotype, rather than to establish the biochemical mechanism underlying the defect.

We hypothesized that impaired βCA activity could limit the conversion of HCO_3_^−^ back to CO_2_, potentially reducing CO_2_ availability at the Rubisco carboxylation site (Badger & Price, 1994). Therefore, we evaluated plant growth under ambient (390 ppm) and elevated (800 ppm) CO_2_ conditions to determine whether increased atmospheric CO_2_ availability could alleviate the mutant phenotypes. As expected, plant height increased significantly in both M82 and *ca1-cr#5* under elevated CO_2_, whereas *ca4-cr#1* showed no significant improvement (Figure 5a and b). Interestingly, leaflet area exhibited a contrasting response: elevated CO_2_ significantly reduced leaflet area in M82 and *ca1-cr#5*, while significantly increasing it in *ca4-cr#1* (Figure 5c). Nevertheless, the leaflet area of *ca4-cr#1* remained significantly smaller than that of the other genotypes under elevated CO_2_. This partial recovery is consistent with previous reports of CO_2_-mediated growth improvement in *Arabidopsis* and tobacco mutants deficient in homologous βCA isoforms (Hines *et al*., 2021; Weerasooriya *et al*., 2022, 2026). However, unlike the growth recovery reported in these species, elevated CO_2_ did not significantly improve plant height or root elongation in *ca4-cr#1* (Figure 5b and d). The persistence of severe root-growth defects despite increased CO_2_ availability suggests that reduced carbon assimilation alone may not fully explain the developmental abnormalities associated with loss of *SlβCA4*.

One possible explanation for the differences in growth recovery between our study and previous reports is the substantially different CO_2_ concentrations applied. Previous studies demonstrated pronounced phenotypic recovery under extremely high CO_2_ concentrations (up to 30,000 ppm), whereas our experiments were conducted at a more moderate concentration of 800 ppm. Thus, although 800 ppm CO_2_ was sufficient to partially improve leaflet expansion in *ca4-cr#1*, higher CO_2_ concentrations may be necessary to alleviate its more severe developmental defects. Whether such concentrations could restore root growth and overall plant development remains to be determined. Together, these findings indicate that elevated CO_2_ can partially alleviate specific aspects of the *ca4-cr#1* growth phenotype, while the limited recovery observed at 800 ppm suggests that SlβCA4 may also contribute to physiological or developmental processes that are not readily compensated for by moderate CO_2_ enrichment.

Following our observations of CO_2_-mediated growth responses, we sought to explore the potential involvement of these isoforms in stomatal development, as stomata serve as the primary gateway for CO_2_ influx (Lawson & Blatt, 2014; Engineer *et al*., 2014). Previous studies have established a strong link between fluctuating environmental conditions, including both CO_2_ concentrations and temperature, and stomatal development in *Arabidopsis* and tomato (Engineer *et al*., 2014; Nir *et al*., 2026). Therefore, we assessed the impact of an environmental shift on stomatal development by measuring the stomatal index (SI) under elevated temperature. When the ambient temperature increased from 26°C to 34°C, the control M82 plants exhibited a significant increase in SI, consistent with prior reports in tomato (Nir *et al*., 2026). However, under the exact same temperature manipulation, both the *ca1#5* and *ca4#1* mutants remained entirely unresponsive, displaying no statistically significant changes in their SI (Figure 6a and b). This lack of phenotypic plasticity suggests that both SlβCA1 and SlβCA4 are likely involved in sensing environmental cues and transducing signals required for stomatal developmental pathways. Overall, these findings parallel studies in *Arabidopsis*, which demonstrate that specific βCA isoforms, such as AtβCA1 and AtβCA4, play crucial roles in environmental sensing and the modulation of stomatal development through carbon signaling networks (Hu *et al*., 2010; Engineer *et al*., 2014).

## Conclusions

In conclusion, our characterization of tomato SlβCA1 and SlβCA4 suggests distinct physiological roles for these two chloroplast-localized β-carbonic anhydrases. The severe developmental defects and lack of stomatal responsiveness to temperature shifts observed in *slca4* mutants indicate that SlβCA4 likely plays a critical role in early plant development and environmental sensing. Furthermore, given its putative role in catalyzing the conversion of HCO_3_^−^ to CO_2_, the severe phenotype of *ca4* mutants may stem, at least in part, from a diminished CO_2_ supply to Rubisco during photosynthesis (Badger & Price, 1994). This is supported by our observation that elevated CO_2_ could induce a slight vegetative improvement, suggesting an underlying limitation in carbon availability.

In contrast, the normal growth of *ca1* mutants suggests that other CA isoforms likely compensate for its absence in basic carbon metabolism. Although SlβCA1 localizes to the photosynthetic mesophyll cells, its primary function may not be required for everyday baseline growth. Instead, as indicated by prior studies linking SlβCA1 and related isoforms to basal immunity and pathogen defense (Hu *et al*., 2021; Zhao *et al*., 2023), this isoform might be specialized for stress responses. Therefore, while it may not be essential for standard development, SlβCA1 could play a vital role under specific environmental or biological stresses (Zhou *et al*., 2020). Further research is necessary to fully explain the likely stress-responsive nature of SlβCA1 alongside the critical developmental and photosynthetic role of SlβCA4, which will ultimately expand our understanding of carbon management in crops.

## Acknowledgments

This work was supported by the Ministry of Innovation, Science and Technology, Israel, through the Food Security (FoodTech) 2023 program (research grant no. 1001704046).

## Author Contributions

OM performed research, analysed data, and wrote the paper; MST performed research, analysed data, and wrote the paper; JS, YN, IB, AY, and YK performed research; YZ analyzed data and wrote the paper; IN designed and performed research, analyzed data, wrote the manuscript and supervised the research.

## Data Availability Statement

*Data required for conclusions in this paper is all provided in main text and supplemental materials*.

## Supporting Information

Additional Supporting Information may be found in the online version of this article (see also the accompanying Supporting Information file).

**Fig S1.** Severe developmental defects in independent SlβCA4 CRISPR mutants.

**Fig S2.** Establishment of SlβCA4 mutant shoots grafted onto wild-type rootstocks.

**Fig S3.** Wild-type rootstocks partially improve the growth of SlβCA4 mutant shoots.

**Table S1.** TargetP-2.0 platform prediction output from protein sequences showing mitochondrial localization of SlβCA4, chloroplastic localization of SlβCA1, and non-chloroplastic localization of SlβCA1 CRISPR mutant ca1-cr#5.

**Table S2.** Primers and sgRNAs used in this study.

