## Supplemental figures and tables for "β-Carbonic anhydrase SlβCA4 is required for normal growth, carbon assimilation and stomatal plasticity in tomato"

**This file includes:**

Figures S1–S3
Tables S1–S2

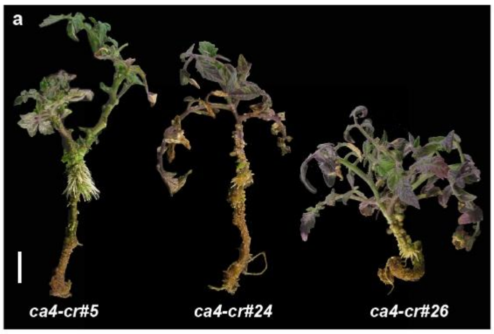

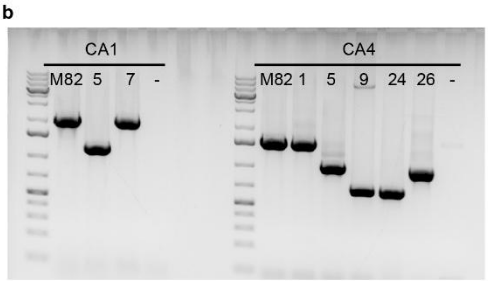

**Figure S1. Severe developmental defects in independent SlβCA4 CRISPR mutants.** (a) Representative T0 plants from three independent homozygous SlβCA4 CRISPR lines, ca4-cr#5, ca4-cr#24, and ca4-cr#26, showing severe developmental defects, including strongly reduced shoot and root growth. Scale bar 2 cm. (b) PCR-based genotyping of SlβCA1 and SlβCA4 CRISPR lines. Amplification products from M82 and the indicated mutant lines demonstrate large genomic deletions at the targeted loci. The SlβCA4 lines shown in (a) carried large deletions and exhibited severe growth defects. "–", negative control; left lane, DNA size marker.

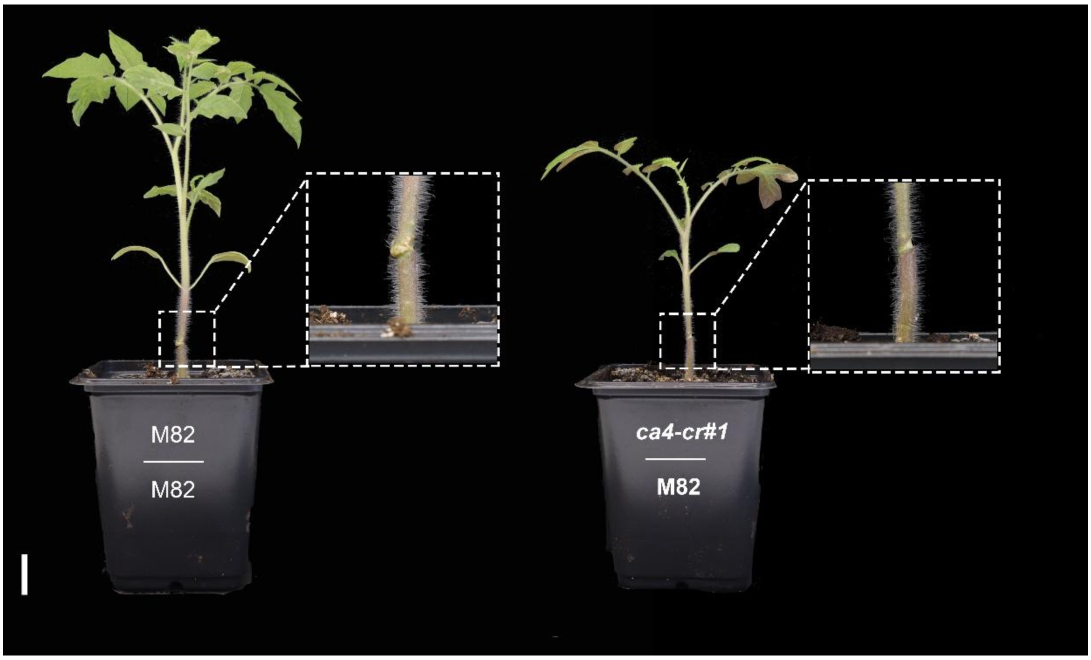

**Figure S2. Establishment of SlβCA4 mutant shoots grafted onto wild-type rootstocks.** Representative M82 self-grafted control (M82/M82) and ca4-cr#1 shoot grafted onto an M82 rootstock (ca4-cr#1/M82) following recovery from grafting. Insets show magnified views of the graft junctions, demonstrating successful graft union formation. Scale bar 2 cm.

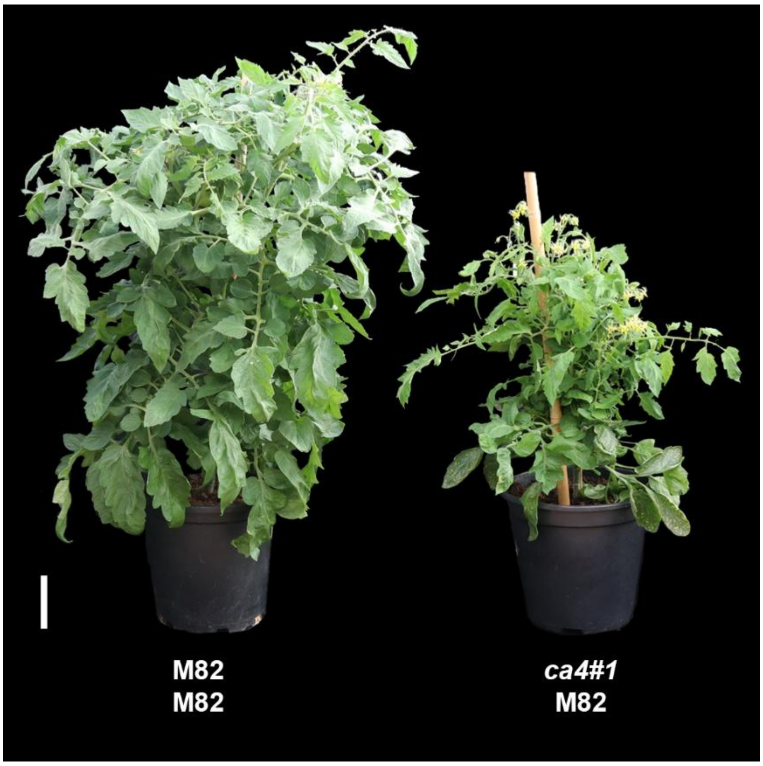

**Figure S3. Wild-type rootstocks partially improve the growth of SlβCA4 mutant shoots.** Representative M82 self-grafted control (M82/M82) and ca4-cr#1 shoot grafted onto an M82 rootstock (ca4-cr#1/M82) two months after grafting. The ca4-cr#1 shoot grafted onto an M82 rootstock was able to establish and undergo substantial vegetative growth, although its growth remained reduced compared with the M82 self-grafted control. Scale bar 5 cm.

**Table S1**: TargetP-2.0 platform prediction output from protein sequences showing mitochondrial localization of SlβCA4, chloroplastic localization of SlβCA1, and non-chloroplastic localization of SlβCA1 CRISPR mutant ca1-cr#5.

| | Cleavage site position (AA) | Other | Signal peptide | Mitochondrial transfer peptide | Chloroplast transfer peptide | Thylakoid luminal transfer peptide |
|---|---|---|---|---|---|---|
| SlβCA1 | 46-47 | 0.0075 | 0 | 0.0047 | 0.9865 | 0.0012 |
| slβca1-cr#5 | None | 0.998 | 0.0002 | 0 | 0 | 0 |
| SlβCA4 | 33-34 | 0.4352 | 0 | 0.563 | 0.0016 | 0.0001 |

**Table S2**: **Primers and sgRNAs used in this study.** List of primers used for construct generation and genotyping, and sgRNAs used for CRISPR-Cas9 mutagenesis.

| Primer or gRNA ID | Sequence 5'-3' | Purpose |
|---|---|---|
| SlCA1g_8_F | TGCTTCCATTAACAATTGCCT | Genotyping and sequencing |
| SlCAg180_R | ATGGGGTTGATGATGGGAGT | Genotyping and sequencing |
| SlCA1g-697_F | TAGAGGTGGCAATTAGGTAGAGG | Genotyping and sequencing |
| SlCA1g_1386F | GAAAtGGCAAAAGAATCCTACG | Genotyping and sequencing |
| SlCA1g_1435R | GTTTCTCGAGTGCAGCAATG | Genotyping and sequencing |
| SlCA1g_2064R | GGTTTGTGCTGTTGGTACATAAA | Genotyping and sequencing |
| SlCA1_174uF | TGGATTGAAGCTCTCATTTGC | Genotyping and sequencing |
| SlCA1g_2073R | ATAAGGCTGGGTTTGTGCTG | Genotyping and sequencing |
| SlCA4_281_F | AAGAAGACTTAATGGCAAACAAATCTTACGAGGAG | Genotyping and sequencing |
| SlCA4_1037_F | CTCGCCTTGGATAACACTGC | Genotyping and sequencing |
| SlCA4_1836_R | TGAAGGATCAGTCTGCACCA | Genotyping and sequencing |
| AATG-slca1-cr#5_F | AAGAAGACTTAATGGCAAAAGAATCCTACGATCAAG | Genotyping, cloning and sequencing |
| SlCA1_ TTGC _R | TTGAAGACAACGAagATACGGAAAGAGGAGGAGAAAGAC | Genotyping, cloning and sequencing |
| SlCA1-mid_R | TTGAAGACTTCCTCCACAAGCACTGTGG | Genotyping, cloning and sequencing |
| SlCA1-mid_F | TTGAAGACTTGAGGTATCAAAGGTCTAATGTCACTA | Genotyping, cloning and sequencing |
| CA1-Ex1_TTCG_R | TTGAAGACAACGAaACAATGGGGTTGATGATGGGA | Genotyping, cloning and sequencing |
| AATG-SlCA1_F | AAGAAGACTTAATGTCAACTGCTTCCA | Genotyping, cloning and sequencing |
| AATG-SlCA4_F | AAGAAGACTTAATGATCTTGAGGACCCAAGTGA | Genotyping, cloning and sequencing |
| SlCA4_ TTGC _R | TTGAAGACAACGAagAGTTCCAGAATTCCCGGTTCT | Genotyping, cloning and sequencing |
| CA1 gRNA2 | GCACATACGAGGATACCCGT | Golden Gate cloning |
| CA1 gRNA3 | GCTGAATTGAAATCAGCAGA | Golden Gate cloning |
| CA1 gRNA5 | GGGACGAGTAGGTTTCTTAA | Golden Gate cloning |
| CA4 gRNA5 | AATTATTCCCGTATTCAACA | Golden Gate cloning |
| CA4 gRNA6 | AATGACCCTGACCAAGGAAG | Golden Gate cloning |
| CA4 gRNA16 | ATATACAACACCTCATAAGG | Golden Gate cloning |
